# A Generalizable Scaffold-Based Approach for Structure Determination of RNAs by Cryo-EM

**DOI:** 10.1101/2023.07.06.547879

**Authors:** Conner J. Langeberg, Jeffrey S. Kieft

## Abstract

Single-particle cryo-electron microscopy (cryo-EM) can reveal the structures of large and often dynamic molecules, but smaller biomolecules remain challenging targets due to their intrinsic low signal to noise ratio. Methods to resolve small proteins have been applied but development of similar approaches for small structured RNA elements have lagged. Here, we present a scaffold-based approach that we used to recover maps of sub-25 kDa RNA domains to 4.5 - 5.0 Å. While lacking the detail of true high-resolution maps, these are suitable for model building and preliminary structure determination. We demonstrate this method faithfully recovers the structure of several RNA elements of known structure, and it promises to be generalized to other RNAs without disturbing their native fold. This approach may streamline the sample preparation process and reduce the optimization required for data collection. This first-generation scaffold approach provides a system for RNA structure determination by cryo-EM and lays the groundwork for further scaffold optimization to achieve higher resolution.

## INTRODUCTION

RNAs often require complex tertiary or quaternary folds to affect a phenotypic response. Understanding the structure-function relationship of an RNA molecule therefore requires a detailed understanding of its three-dimensional shape, at an atomic level if possible. Historically this has been achieved through the use of X-ray crystallography and nuclear magnetic resonance (NMR) spectroscopy. Although powerful, both methods present limitations when working with RNA: X-ray crystallography requires crystallization, and NMR has a practical upper size limit beyond which spectral crowding makes structure determination challenging. Lower-resolution methods such as RNA chemical probing and small angle X-ray scattering (SAXS) supplement our structural understanding of RNA, but do not provide sufficient information to reliably and routinely provide unambiguous three-dimensional RNA structural information^1–4^.

Single-particle cryo-electron microscopy (cryo-EM) now allows determination of atomic resolution structures of many biomolecules and biomolecular complexes^5–7^, including those with intrinsic conformational dynamics^8, 9^. However, this has largely been limited to protein and larger protein-nucleic acid complexes, with relatively few RNA-only structures determined by cryo-EM^10^. This can, in part, be explained by conformational dynamics in many RNAs, which complicates structural studies^11–13^, and because many RNA targets of interest are <50 kDa in mass^14–16^. The small size of such RNAs results in a low signal to noise ratio in the micrographs, making high-resolution structure determination by cryo-EM difficult or intractable^17, 18^. Indeed, while improved cryo-EM hardware and sample preparation makes it increasingly possible to achieve higher resolutions for smaller targets^19^, a theoretical lower limit on single-particle cryo-EM as applied to proteins was proposed as 38 kDa^20^. A number of properties of RNA theoretically may make it more amenable to single particle cryo-EM, such as the presence of phosphorus in the backbone, resulting in greater contrast with the solvent compared to protein. Additionally, RNA’s lower density compared to protein results in larger 2D projections for the same mass. However, these gains in signal to noise may be offset by RNAs dynamic nature, in part a result of the sparse tertiary interactions which stabilize the overall fold of many RNA elements. Hence, the theoretically lower limit of RNA is likely not too dissimilar to that of proteins. When it is possible to recover a cryo-EM map of very low molecular weight RNA molecules, model building is complicated by low resolution and difficulty in determining the base register and stem identity, often requiring alternative structure modeling approaches^21–24^.

To circumvent the current size limitation of cryoEM, various approaches have been developed for protein cryo-EM. These include the use of fragment antigen-binding regions (FAbs) to increase the effective size of a biomolecule^17, 25–27^, and scaffold-based systems that often use highly symmetric protein complexes to display a cargo of interest^25, 28, 29^. However, similar approaches for RNA-only cryo-EM are sparse, despite the fact that RNA elements of interest are often smaller than the theoretical lower limit^14–16^.

We have developed a scaffold-based approach in which an RNA sequence of interest is engineered into a well-ordered larger RNA structure for single-particle cryo-EM structure determination. We first determined that a circularly permuted version of the Tetrahymena ribozyme is an appropriate scaffold, with a rigidity ideal to “display” smaller RNAs. As a proof of principle, we tested this scaffold approach using three RNAs of known structure: (1) a 23 kDa element from Zika Virus, (2) a 17 kDa Fluoride riboswitch, and (3) a 17 kDa element from Tamana Bat Virus. The resultant maps of the appended RNAs had resolutions of 5.04 Å, 4.95 Å, and 4.46 Å respectively, sufficient for *ab initio* structural modeling. While the maps did not allow de novo assignment of bound ligands such as magnesium ions, their presence was visible in the maps. Our results illustrate the potential generalizability of this approach to determine RNA structures which are significantly smaller than the generally accepted lower cryo-EM size limit. Even when high-resolution maps are not obtained, information about the architecture and behavior of folded RNAs can be assessed.

## RESULTS

### Motivation and design considerations for a scaffold-based approach

We reasoned that we could take advantage of the modularity of RNA structure^30^, which is largely composed of A-form helices connected by linker regions and whose 3-D fold is stabilized by tertiary interactions^31^. It is often possible to alter distal regions of RNA stem-loops extended away from a core central fold without compromising the overall folded structure or function of the RNA. Indeed, this is a proven method to obtain RNA crystals^32^. Appending a small RNA of interest onto an available stem-loop of a larger, well-ordered, recognizable, and robustly folded scaffold would increase the effective size of the molecule, making it more amenable to cryo-EM^20^ (Fig.1A). Also, this strategy would ensure a 1:1 stoichiometry, help enforce proper folding, allow transcription *in vitro*, and prevent the need to optimize binding conditions necessary with methods that use intermolecular interactions^17, 26, 27, 33^. In addition, if the RNA domain is engineered onto the scaffold molecule, the helical orientation and the register of the RNA are known, simplifying model building and removing ambiguity. Other methods to address this often require collecting data on several constructs and determining helical orientation by process of elimination^23^. Finally, some RNAs are difficult to characterize structurally due to intrinsic dynamics and heterogeneity^13, 23^. By applying this approach to RNAs with the propensity to misfold and aggregate, we might mitigate both to yield usable cryo-EM samples.

**Figure 1.**
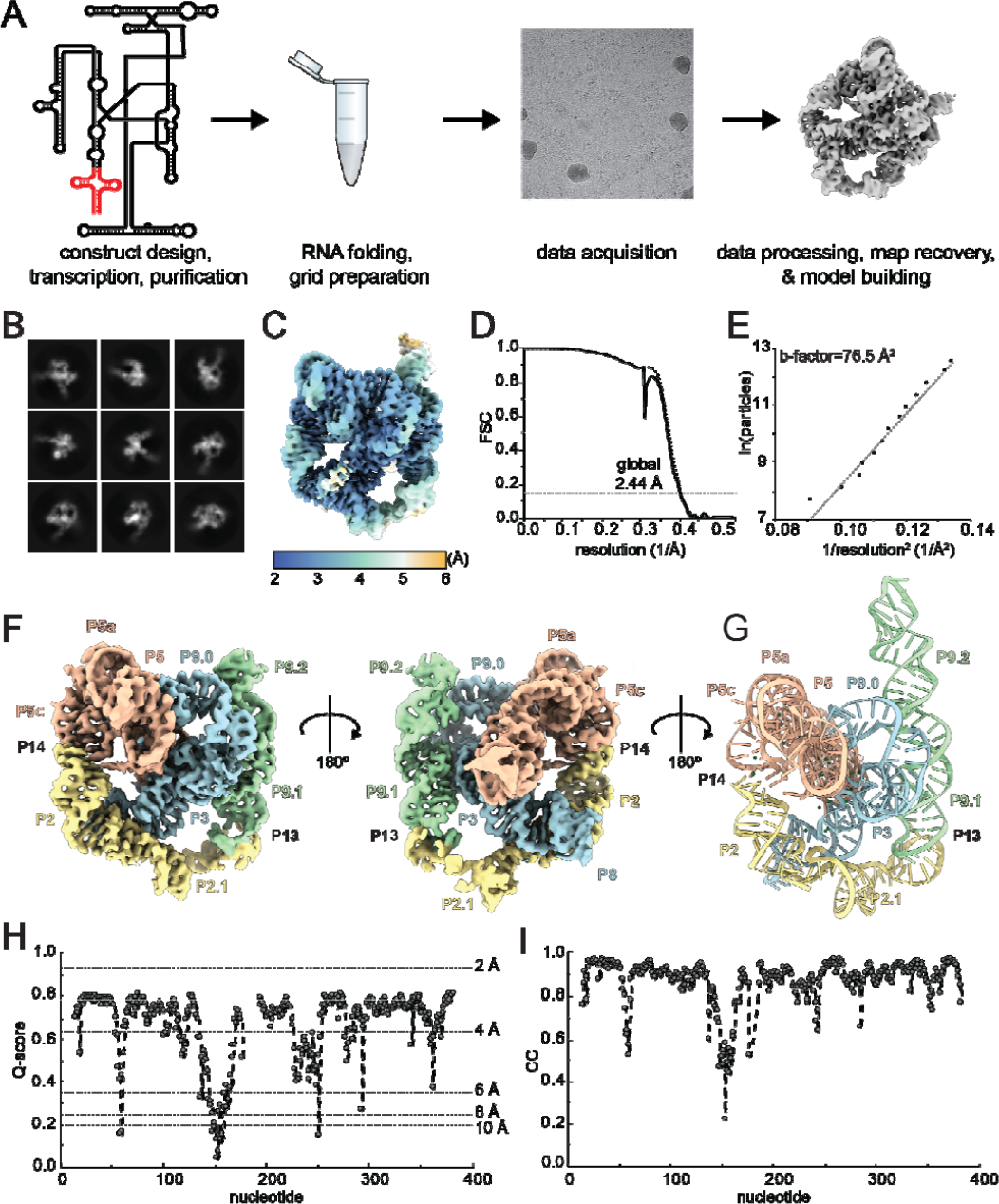
Workflow of the scaffold-based approach and identification of a suitable scaffold. **A.** The construct has the RNA of interest (red) appended to the larger group I intron RNA. After transcription and purification, the pure RNA is subjected to a standard RNA folding protocol, applied to cryo-EM grids and vitrified. Data collection and processing yields a map for model building and refinement. **B.** Representative 2D class averages of the Tet_P6b_ construct. **C.** Local resolution map of the recovered Tet_P6b_ construct. **D.** Fourier Shell correlation graph of the Tet_P6b_ construct with tight mask (dashed line) and noise subtracted (solid line). **E.** Henderson-Rosenthal plot of the Tet_P6b_ construct. **F.** Recovered map of the Tet_P6b_ construct. **G.** Resulting atomic model of the Tet_P6b_ construct. **H.** Per-nucleotide Q-score graph of the Tet_P6b_ construct. **I.** Map-to-model cross-correlation score graph of the Tet_P6b_ construct.

### Tet_P6b_ is an ideal scaffold for high-resolution structure determination

Decades of work have characterized the Tetrahymena group I intron RNA’s structure and conformational dynamics^34–36^, revealing features that suggest it is a good scaffold for our approach. To test the amenability of this RNA, used a circularly permuted version (hereafter referred to as Tet_P6b_). The circularly permuted version of the Tetrahymena ribozyme has the normal 5’ and 3’ ends connected, giving rise to 5’ and 3’ ends in a new position in the structure. In this case, the 5’ and 3’ ends were moved to the P6b helix loop. We prepared grids with this RNAs, then collected 10,422 movie images using a Titan Krios equipped with a BioQuantum K3 energy filter. Following template-based picking, we performed several rounds of rigorous two-dimensional (2D) classification, removing poor quality 2D classes and the corresponding particles (Extended Data Fig. 1). The good classes clearly correspond to projections of the RNA with visible structural features such as major and minor grooves (Fig. 1B). These particles were used for initial map determination followed by cryoSPARC’s non-uniform refinement^37^ and local motion correction and CTF refinement steps. The resulting map of the Tet_P6b_ yielded a nominal global resolution of 2.44 Å (Fig. 1C,D). To date, this is one of two published RNA-only cryo-EM maps below 2.5 Å resolution^38^. In most regions of the map, the sugar-phosphate chain is clearly resolved, allowing for unambiguous tracing of the entire RNA backbone with sufficient detail to allow *de novo* building of the structure (Fig. 1F,G & Extended Data Fig. 2). In some parts of the map (mainly around the core domains P7-P3-P8 and most of P4-P6), nucleobases are clearly resolved with features such as exocyclic amines and carbonyl oxygens visible (Extended Data Fig. 2). This resolution allowed unambiguous placement of non-Watson-Crick base interactions and native magnesium ions, whose identity was previous known (Extended Data Fig. 2).

We assessed the quality of the map and model by calculating a per-nucleotide Q-score to determine the local quality of the map^39^, and map to model cross-correlation (CC) score to ensure the built model was in agreement with recovered map^40^ (Fig. 1H,I). The Q-score remained within a range expected for a map with a resolution range of 2-4 Å, and the CC score was comparably high. Dips in both the Q-score and the CC score matched the lower quality regions of the map such as peripheral helices and loops, as well as regions known to be more dynamic or structurally heterogenous such as the kissing loop P13. These trends were also observed in prior cryo-EM studies of the Tetrahymena ribozyme^33, 41, 42^.

Because our method is designed to recover high-resolution maps of biomolecules, ideally the use of additional particles would result in an even higher-resolution map. Indeed, a Rosenthal-Henderson plot shows a high correlation between particle number and resolution (Fig. 1E). Thus, additional particles should contribute to increased resolution and resolution is therefore not limited by the intrinsic dynamics of the RNA nor by technical limitations. We did not choose to pursue higher resolution of this RNA, as our goal was to identify a suitable scaffold for displaying smaller RNAs; this analysis achieved that.

This is the first example of a cryo-EM structure of a circularly permuted version of the Tetrahymena group I intron. In this precatalytic state, the ΩG is poised for the first transesterification step in splicing, surrounded by several well-ordered magnesium ions (Extended Data Fig. 2). Unsurprisingly, the active site of the circularly permuted structure remains largely the same as in the solved structure of the wild type RNA (Extended Data Fig. 3). However, the regions corresponding to P1 and the 3’ end of P9.0 that comprise the linker are largely unstructured in the circularly permuted version of the ribozyme, making it difficult to trace the RNA here (Extended Data Fig. 3). Importantly for the development of our method, this analysis shows Tet_P6b_ is a good candidate for a scaffold-based approach. Notably, circularly permuting and appending domains onto stem-loops other than P6b did not yield usable data, hence we focused on Tet_P6b_ as the scaffold candidate.

### The structure of a 23 kDa RNA element from Zika Virus recovered by cryo-EM

To benchmark this scaffold-based method, we tested it with three RNAs of known structure solved by X-ray crystallography. The first was a compact exoribonuclease-resistant RNA (xrRNA) from Zika Virus^40^ (Fig. 2). We generated a construct with the P2 stem of this 23 kDa RNA fused to the P6b stem of Tet_P6b_. Importantly, the xrRNA’s P2 stem is not directly involved in the functional motif of the xrRNA and thus was ‘available’ to be appended to the scaffold^43^. Following transcription of this construct, we analyzed the sample using nondenaturing (native) gels; these were consistent with the RNA folding into a single state and suggested the sample was amenable to single particle cryo-EM analysis (Extended Data Fig. 4).

**Figure 2.**
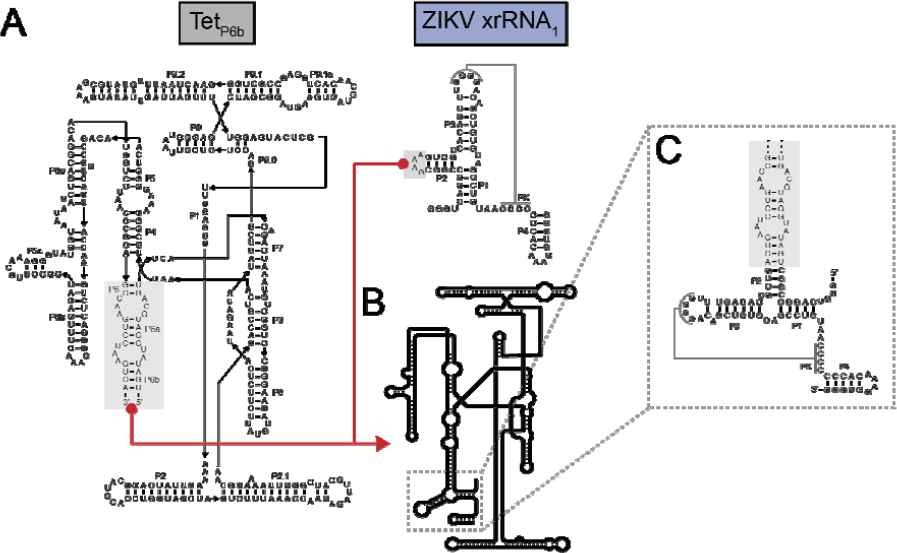
Engineering of the Zika Virus xrRNA-containing test RNA. **A.** Secondary structures of the Tet_P6b_ RNA and the Zika Virus xrRNA (shaded blue) used in this study. The two RNAs were appended at the locations boxed in grey and designated with red lines. In the xrRNA, P2 can be altered without affecting xrRNA function, therefore the apical loop of this stem was removed and became the point of attachment. **B.** Cartoon secondary structure depictions of the chimeric RNA, with the xrRNA in blue. **C.** Close-up of the engineered region.

We collected 7030 movie images using a Titan Krios equipped with a BioQuantum K3 energy filter (Extended Data Fig. 5). Following motion correction and CTF estimation in cryoSPARC, particles were picked with templates generated from the first 1000 micrographs. After several rounds of rigorous 2D classification, the appended Zika xrRNA was observable (Fig. 3A). Using the resulting 389,185 particles, initial map generation yielded a class where the appended xrRNA was clear (Fig. 3B), but map quality was poor in the distal regions of the xrRNA’s P4 stem. To achieve higher resolution of the xrRNA’s distal regions, we parsed this pruned dataset with three-dimensional (3D) classification approaches, revealing some conformational heterogeneity in the distal P4 stem, likely reflecting local inherent motion (Extended Data Fig. 6). To best resolve this region, we only proceeded with 3D classes in which the maps were strong for this entire region. The resulting 3D class of 68,847 particles was used for final map refinement, yielding a nominal resolution of the total RNA of 3.40 Å (Fig. 3C,D), with the highest resolution in the Tet_P6b_ scaffold. Local resolution assessment showed a nominal resolution of 5.04 Å for the xrRNA domain, with decreasing resolution in the distal regions, largely in P4 (Fig. 3C,D).

**Figure 3.**
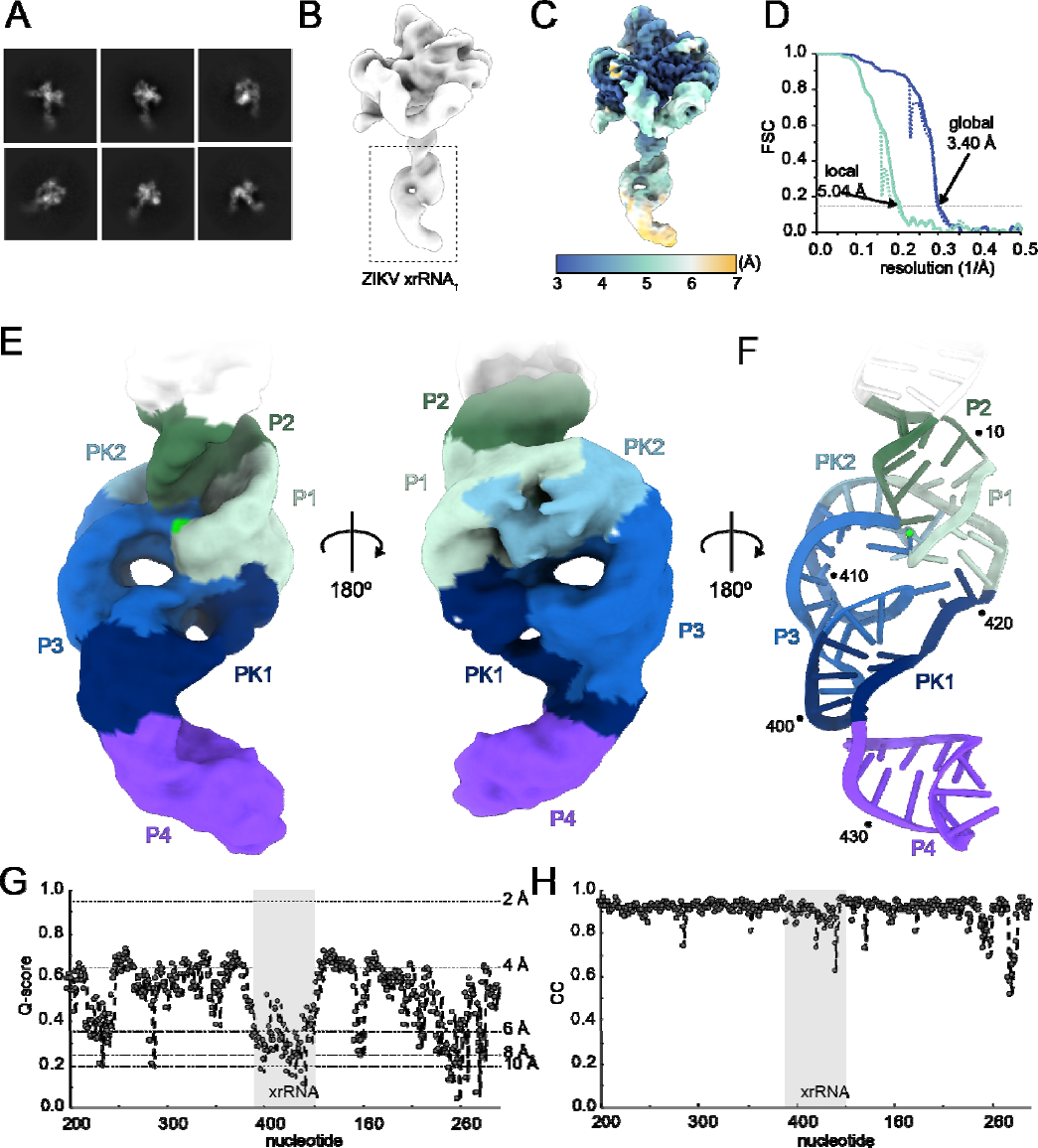
Cryo-EM map of the Zika Virus xrRNA appended onto a scaffold. **A.** Representative 2D class averages of the Zika Virus xrRNA+Tet_P6b_. **B.** Cryo-EM map indicating the location of the Zika Virus xrRNA in the overall initial map. **C.** Local resolution map of the chimeric Zika Virus xrRNA construct. **D.** Fourier Shell correlation graph of the Zika Virus xrRNA construct with global resolution (blue) and the local xrRNA resolution (teal) using a tight mask (dashed line) and noise subtracted (solid line). **E.** Recovered map of the chimeric Zika Virus xrRNA construct. **F.** Resulting atomic model of the chimeric Zika Virus xrRNA construct. A visible magnesium ion is shown as a bright green sphere. **G.** Per-nucleotide Q-score graph of the Tet_P6b_ construct. **H.** Map-to-model cross-correlation score graph of the Tet_P6b_ construct.

Clear structural features were apparent within the part of the map that contained the appended xrRNA, such as unambiguous major and minor grooves in A-form regions (Fig. 3E). Additionally, while individual bases were not directly resolved, clear bumps in the maps for the phosphates of the sugar-phosphate backbone in regions proximal to the scaffold allowed for unambiguous tracing of the backbone. Remarkably, the distinctive ring-like architecture, a defining feature of xrRNA structures determined to date, was clearly visible, and the map showed features consistent with a magnesium ion at the core of the fold that was seen in two crystal structures. The limited resolution in this region was not sufficient to place this ion *de novo*, but was sufficient to confirm the presence of the ion, lend support for its structural importance, and confirm it was not an artifact of crystallization. Hence, globally, the map matches the known structure and confirms many important features.

Using the scaffold to ensure the correct register, we built an *ab initio* atomic model of the Zika xrRNA, relying on the known secondary structure of the xrRNA (Fig. 3F). We then evaluated the map and model with the Q-score^39^ and map-to-model CC score^40^ (Fig. 3G,H). The map of the xrRNA+Tet_P6b_ RNA had Q-scores in line with the observed 3.4 Å resolution through much of Tet_P6b_ and consistently high CC scores, both validating the model. Within the xrRNA regions of the map, Q-values ranged from 0.5 to 0.2, in agreement with the observed local resolution in this part of the map while suggesting the model was not overfit.

Overall, the Zika Virus xrRNA structural model obtained from cryo-EM using our scaffold approach agreed well with the high-resolution crystal structure of the Zika xrRNA^43^, with an average backbone RMSD of 3.0 Å across the entire structure and 2.1 Å when P4 is excluded (Fig. 4A,B). This is largely because the angle between the P4 stem relative to the rest of the structure was shifted 14° compared to its position in the crystal structure (Fig. 4A). Interestingly, in the crystal structure of another xrRNA (from Murray Valley Encephalitis Virus), the P4 stem is nearly perpendicular to its position in the Zika xrRNA structure^44^ (Extended Data Fig. 7). The cryo-EM map data now support the idea that the P4 stem is mobile compared to the rest of the molecule, and different positions were captured in the two crystal structures. The functional significance of this motion is outside the scope of this study. However, as cryoEM allows visualization of molecular motions often precluded by crystallization suggests this scaffold-based approach may be useful for detecting or confirming putative dynamic regions that can then be studied with other methods.

**Figure 4.**
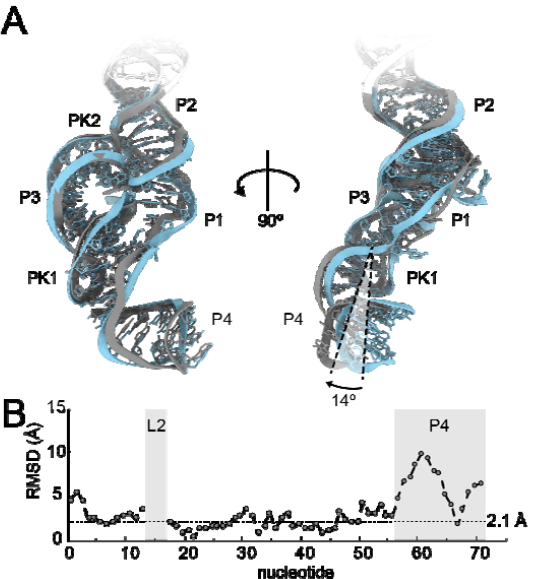
Comparison of cryo-EM-derived model with the X-ray crystallography structure of the Zika Virus xrRNA. **A.** Overlay of the cryo-EM-based model of the Zika Virus xrRNA (grey) and the crystallography-based model of the Zika Virus xrRNA (cyan). **B.** Per-nucleotide backbone RMSD graph of both Zika Virus xrRNA structures. The RMSD of the L2 loop is omitted as the analogous nucleotides are not present in the cryo-EM model.

In summary, while we did not achieve true atomic resolution in the xrRNA, the quality of the map was sufficient to reveal key architectural features, potential conformational dynamics, and for building and refining a biologically informative structural model. Thus, the scaffold approach is a viable method for mid-resolution structure determination of RNAs smaller than the traditional lower size limit of cryo-EM. When combined with orthogonal data such as covariation analysis and chemical probing, the approach may provide sufficient structural information to build high-confidence models of RNAs difficult to resolve by other high-resolution structural techniques. In addition, given that this structure was obtained with only ∼68K particles, addition of more data may lead to maps of increased resolution.

### Testing the method on two additional small (<20 kDa) RNAs

We tested the general applicability of our scaffold method, applying it to two additional examples: (1) a ∼17 kDa xrRNA from Tamana Bat Virus (TABV)^45^, distinct in sequence from the Zika Virus xrRNA, and (2) a ∼17 kDa Fluoride riboswitch from *Thermotoga petrophila*^46^ (Fig. 5). Similar to the scaffolded Zika xrRNA, we used native gels to qualitatively determine that these RNAs were amenable to single particle cryo-EM (Extended Data Fig. 4). Then, using the same basic workflow as for the Zika xrRNA construct, we obtained the expected architecture of the scaffold (Extended Data Fig. 8,9). Both resulting maps clearly contain the appended domains with visible structural features such as major and minor grooves.

**Figure 5.**
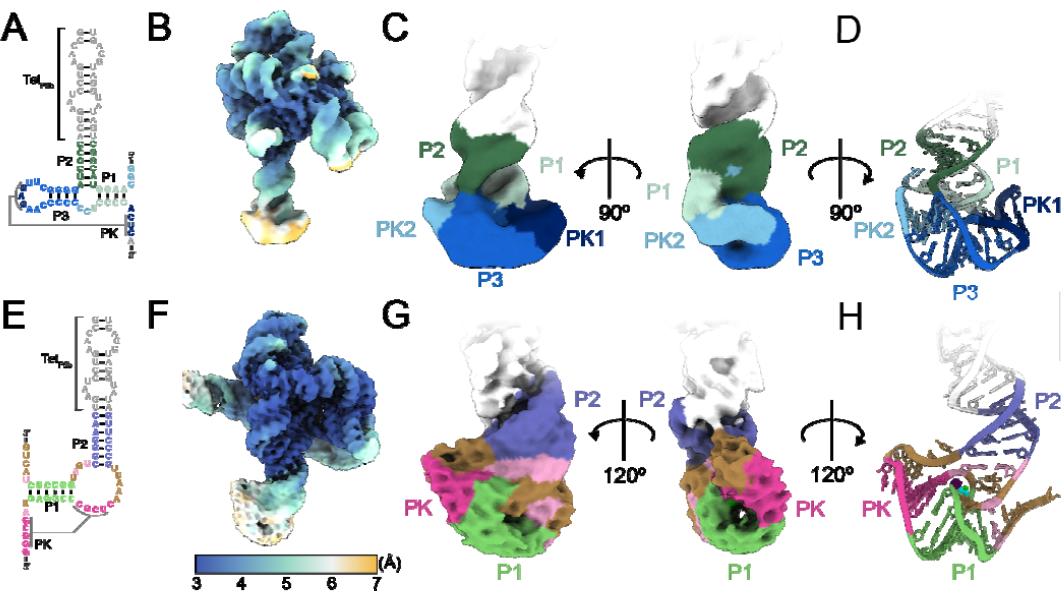
Additional examples of two ∼17 kDa RNA structures recovered using cryo-EM. **A.** Secondary structure depiction of the TABV xrRNA appended onto Tet_P6b_. **B.** Local resolution map of the recovered chimeric TABV xrRNA construct. **C.** Recovered map of the chimeric TABV xrRNA construct. **D.** Resulting atomic model of the chimeric Zika Virus xrRNA construct. **E.** Secondary structure depiction of the chimeric *Thermotoga petrophila* Fluoride riboswitch appended onto Tet_P6b._ **F.** Local resolution map of the Fluoride riboswitch construct. **G.** Recovered map of the chimeric Fluoride riboswitch construct. **H.** Resulting model of the chimeric Fluoride riboswitch construct.

Within the map of the TABV xrRNA at 4.95 Å, the backbone of the RNA is traceable through the entirety of the structure, with clear phosphate bumps visible in some parts of the map, specifically in regions proximal to the scaffold domain (Fig. 5A-C). Similar to the map of the Zika xrRNA, while specific base interactions could not be built *de novo* through the entirety of the structure, the map provides sufficient structural information such that when combined with orthogonal RNA structural techniques such as chemical probing and sequence alignments, we could build much of the structure without relying on the solved X-ray crystallography structure (Fig. 5D). The resulting model demonstrated backbone RMSD of 1.5 Å compared to the crystal structure throughout the structure, again validating this approach as a viable method to model RNA structures.

The map of the Fluoride riboswitch at 4.46 Å showed similar structural features as the TABV xrRNA, though the quality of the resulting map made data analysis and structure modeling more complicated (Fig. 5E-G). However, structural features distinct to this RNA element were clearly present, including density coorresponding to a pseudoknot that does not lie in the core of the riboswitch (Fig. 5H). Similar to the TABV example, the backbone RMSD compared to the crystal structure throughout the appended domain was calculated at 1.9 Å.

These two additional examples suggests that this scaffold method is generalizable and provides a benchmark for its use. Although high-resolution maps of smaller RNA elements are unlikely to be routinely obtained using this method, the ease of appending different RNA elements of interest allows rapid determination of intermediate-to-high-resolution maps of RNAs for model building, likely without requiring exhaustive optimization of RNA folding and grid preparation.

## DISCUSSION

Single-particle cryo-EM is now a powerful method for determining macromolecular structure. However, the intrinsic signal to noise limits of the data still make analysis of small biomolecules or complexes technically challenging^17, 18^. Whereas methods to circumvent this have been established for protein^17, 25–27^, similar approaches have not been readily available for RNA, which is particularly limiting as many classes of functional RNAs are small for traditional single-particle cryo-EM^14–16^. Here, we demonstrate a method that can help address this limitation, leveraging RNA’s intrinsic modular nature to append diverse RNA elements onto a large scaffold.

This ‘first generation’ of the method has several desirable features. First, this method consistently yielded interpretable maps of the combined Tet_P6b_ scaffold and the appended RNA structures (Zika xrRNA at 5.04 Å, TABV xrRNA at 4.95 Å, and a Fluoride riboswitch at 4.46 Å as assessed using currently accepted map resolution determination methods). Thus, realtively small RNA elements can be examined at resolutions sufficient to gain insight. Intermediate-resolution structures are likely to allow tracing of the backbone of many RNA molecules, and when combined with results from other techniques, biologically relevant and testable tertiary structure models can result. In addition, in some cases the resolution of the map may allow complete or partial structures to be built *de novo.* In other instances, inherent dynamics in the RNA may give rise to uninterpretable maps, but even in those cases information regarding the dynamic behavior of the RNA can guide future studies. A similar approach has previously been described where a small tetraloop-tetraloop receptor was appended to a small viral frameshifting pseudoknot^47^, however this method yielded a relatively low-resolution map that required extensive computational modeling and may not be generalizable^48^.

The method described here simplifies sample preparation and data analysis, as the same large scaffold is used for every new RNA examined. As most appended RNAs are likely much smaller than the scaffold, we predict similar folding procedures, sample concentration, and grid preparation parameters can be initially used for each new RNA construct, with little or no optimization needed (Fig 1A). Data analysis is simplified as the Tet_P6b_ RNA particles are readily recognized and picked using an established template; we observed no preferred orientation, and the maps of the scaffold are relatively easy to recover with a standard workflow adapted by the user to each dataset.

We chose the Tetrahyamena group I intron as the basis for the scaffold as we and others have found it highly amenable to structure determination by cryo-EM^33, 41, 42^. Indeed, here we recovered a map of the circularly permuted Tetrahymena ribozyme at 2.44 Å, similar to the resolutions obtained for other versions of the ribozyme^33, 41, 42^. Remarkably, there is still room for improvement as this scaffold is still in a regime where additional particles could yield higher resolution in both the maps of the scaffold and the appended domains. This bodes well for future applications of this or similar scaffolds.

Although this method can be applied immediately with many benefits, there are features that can be optimized or improved. First, as expected, we observed that within the maps of the appended RNAs, the regions of the maps more proximal to the scaffold showed clearer features. This suggests that it is best to use the shortest possible helix to attach the appended RNA, as long as steric clash does not interfere with the fold. This could be optimized for many RNAs. Related, if an RNA of interest has several available stem-loops that can be altered without loss of function, more than one construct could be examined and the information from multiple maps combined. However, one important limitation of the approach is that RNA elements that lack an available stem-loop element (e.g. many frameshifting pseudoknots and similar) cannot be appended to the scaffold. Finally, while the scaffold we chose has several highly desirable features, other versions of this scaffold or other larger well-folded RNAs could be tested in ‘second-generation’ versions.

In recent years, interest in RNA 3D structure determination has increased. The nature of RNA structure and the fact that far fewer RNA structures have been solved relative to protein means that prediction of RNA 3-D structures is not yet robust. Machine learning and similar methods hold promise, but require many more RNA structures to examine than currently are available, mandating ways to accelerate experimental RNA structure determination. The application of cryo-EM to RNA structure determination is promising, but thus far has been limited for many technical reasons. The method we present here has the potential to rapidly provide intermediate- and high-resolution structural information on a wide variety of structured RNAs, providing biological insight, testable models, and a richer understanding and database of RNA structure.

## METHODS

### *in vitro* RNA transcription and purification

DNA templates were ordered as gBlock DNA fragments (IDT) and cloned into pUC19. 200 µL PCR reactions using primers containing an upstream T7 promoter were used to generate dsDNA templates for transcription. Typical PCR conditions: 100 ng plasmid DNA, 0.5 µM forward and reverse DNA primers, 500 µM dNTPs, 25 mM TAPS-HCl (pH 9.3), 50 mM KCl, 2 mM MgCl_2_, 1 mM β-mercaptoethanol, and Phusion DNA polymerase (New England BioLabs). dsDNA amplification was confirmed by 1.5% agarose gel electrophoresis. Transcriptions were performed in 1 mL volume using 200 µL of PCR product (∼ 0.1 µM template DNA) and 10 mM NTPs, 75 mM MgCl_2_, 30 mM Tris-HCl (pH 8.0), 10 mM DTT, 0.1% spermidine, 0.1% Triton X-100, and T7 RNA polymerase (generated recombinantly in lab). Reactions were incubated at 37°C overnight. After transcription, insoluble inorganic pyrophosphate was removed by centrifugation at 5000xg for 5 minutes, then the RNA-containing supernatant was precipitated with 3 volumes of 100% ethanol at −80°C for a minimum of 1 hour and then centrifuged at 21000xg for 30 minutes at 4°C to pellet the RNA, and the ethanolic fraction was decanted. The RNA was resuspended in 9 M urea loading buffer then purified by denaturing 10% PAGE. Bands were visualized by UV shadowing then excised. Bands were then crush-soaked in diethylpyrocarbonate-treated (DEPC) milli-Q water at 4°C overnight. The RNA-containing supernatant was concentrated using spin concentrators (Amicon) to the appropriate concentration in DEPC-treated water. RNAs were stored at −80°C with working stocks stored at −20°C.

### RNA refolding

To the RNA solution (usually 18 µL of 3 mg/mL, or approximately 25 µM), 1/20^th^ the volume of 1 M Tris pH 7.5 was added. The solution was heated to 90°C for 3 minutes then allowed to cool to room temperature for 10 minutes. Subsequently, 1/20^th^ the volume of 200 mM MgCl_2_ was added. The solution was heated to 50°C for 30 minutes, then allowed to cool to room temperature for 10 minutes. Once cooled, the folded RNA solution was stored on ice until use. For the Fluoride riboswitch construct, 1/20^th^ the volume of 20 mM NaF was added at the time the MgCl_2_ was added, and all other steps remained the same.

### Native gel analysis

To qualitatively assay the folding of each RNA, samples were subject to native gel analysis. Briefly, 1 µg of RNA was refolded as described above. To this, an equal volume of 2X native loading buffer (100 mM Tris pH 7.5, 20 mM MgCl_2_, 20% sucrose) was added. The samples were then loaded on a 5% native acrylamide gel (5% 37.5:1 acrylamide:bisacrylamide, 0.5x TBE buffer, 10 mM MgCl_2_, 5% sucrose) and run at 100 V at 4°C for 2 hours. RNA was visualized by methylene blue staining of the gel and imaged.

### Cryo-EM sample preparation and data acquisition

Each RNA sample was applied to a glow-discharged C-flat (1.2/1.3, 400 mesh) holey carbon grid. Samples were vitrified in liquid ethane using a Vitrobot Mark IV (5 s wait time, 5.5 s blot, −5 blot force, 100% humidity, 4°C). Samples were imaged on a Titan Krios equipped with a Gatan K3 camera and a Bioquantum energy filter. Movies were collected in fringe-free mode, with a physical pixel size of 0.8464 Å, total dose of 50 e^-^/Å^2^, and a defocus range of −0.8 to −2.0 µm. SerialEM was used for all data collection. Additional information on data collection can be found in Supplemental Table 1.

### Cryo-EM data processing

The general data processing workflow was approximately equivalent for all datasets using cryoSPARC 4.0. Briefly, all data were patch motion corrected and the contrast transfer function was locally fit. An initial set of particles was picked using blob picker from the first 1000 micrographs. Particles were extracted and 2D classified. Classes which displayed RNA-like features were were manually chosen for template picking. Template-based picking was performed using template picker on the entire dataset. Particles were then extracted and subjected to several rounds of 2D classification to sort out junk particles (i.e. particles which did not coorrespond to the consensus structure). Good 2D classes, which demonstrated features of RNA such as discernable A-form helices, and the associated particles were manually curated after each round. The pruned particles were then used for initial map generation using the *ab initio* reconstruction job requesting 2 volumes. The resulting 2 volumes were used for heterogenous refinement, where one class was the “good” class and the other a “junk” class to remove poor quality particles. Good classes were those which demonstrated a map in line with the expected scaffold domain, whereas bad classes did not demonstrate RNA-like features. Several rounds of heterogenous refinement were performed, discarding the “junk” particles (i.e. those which did not match the consensus model) after each round until the map quality stopped improving. Subsequently, the “good” volume and particles were subjected to non-uniform refinement. Local motion correction, local CTF refinement, and global CTF refinement were then carried out on the data and the particles were subjected to further rounds of nonuniform refinement to yield the final maps. Additional information on data processing for each map can be found in Extended Data Figures 1, 5, 8, and 9.

### Structure modeling

To build each structure, the cryo-EM structure of the Tetrahymena ribozyme was placed into the maps using rigid body fitting (PDB:7EZ2) followed by manual building of the appended structure. In some regions the solved crystal structure of either the TABV xrRNA (PDB:7K16) or the Fluoride riboswitch (PDB:4ENC) was used to supplement manual building. The structures were then manually manipulated and joined in COOT. The final model was refined against the map with several rounds of real-space refinement in PHENIX and manual adjustments in COOT. The statistics of model refinement and validation are listed in supplemental table 1.

## ACKNOWLEDGEMENTS

The authors thank current members of the Kieft Lab for critical reading of the manuscript and insightful comments. Theo Humphries (PNCC) assisted with microscope operations. This work was supported by NIH grants R35GM118070 and R01AI133348 to J.S.K. A portion of this research was supported by NIH grant U24GM129547 and performed at the PNCC at OHSU and accessed through EMSL (grid.436923.9), a DOE Office of Science User Facility sponsored by the Office of Biological and Environmental Research.

## AUTHOR CONTRIBUTIONS

C.J.L. and J.S.K. conceptualized and designed the research. C.J.L. prepared the RNA samples for cryo-EM. C.J.L. analyzed the cryo-EM data. C.J.L. performed modeling building for all RNAs herein. C.J.L. and J.S.K. wrote the manuscript.

