## Extended Data Figures for "A Generalizable Scaffold-Based Approach for Structure Determination of RNAs by Cryo-EM"

**Contents:**

**Extended Data Figure 1:** Cryo-EM structure determination of Tet_P6b_

**Extended Data Figure 2:** Tet_P6b_ reconstruction details

**Extended Data Figure 3:** Comparison of Tet_P6b_ to a published Tetrahymena ribozyme structure

**Extended Data Figure 4:** Native gel analysis of Scaffolded RNAs

**Extended Data Figure 5:** Cryo-EM structure determination of Tet_P6b_ + Zika Virus xrRNA

**Extended Data Figure 6:** Tet_P6b_ + Zika Virus xrRNA principal component analysis

**Extended Data Figure 7:** Comparison of Tet_P6b_ + Zika Virus xrRNA with crystal structures of other xrRNAs

**Extended Data Figure 8:** Cryo-EM structure determination of Tet_P6b_ + TABV xrRNA

**Extended Data Figure 9:** Cryo-EM structure determination of Tet_P6b_ + *T. petrophila* Fluoride riboswitch

**Extended Data Figure 1. Cryo-EM structure determination of Tet_P6b_.** The Cryo-EM processing pipeline in cryoSPARC for the Tet_P6b_ construct.

**Extended Data Figure 2. Tet_P6b_ reconstruction details. A.** Representative maps and placed nucleobases showing exocyclic amines (blue asterisks) and carbonyl (red asterisk). **B**. Model of the Tet_P6b_ RNA colored by domain. **C.** Density and model of the P7 stem. **D**. Contacts made by the ΩG in the active site. **E.** Kissing loop interaction between P5c and P2 which form P14 with native magnesium ions. **F.** Non-Watson-Crick base triple in P14. **G.** Density and model of the P3 stem. **H.** Long-range P7-P9 interaction.

**Extended Data Figure 3. Comparison of Tet_P6b_ to the published tetrahymena ribozyme structure.** **A.** Map and model of the published tetrahymena ribozyme and the circularly permuted version of the tetrahymena ribozyme from this study. **B.** RNA backbone cross-correlation between the wild type and circularly permuted tetrahymena ribozyme, numbered by the wild type sequence.

**Extended Data Figure 4. Native gel analysis of scaffolded RNAs.** A native gel of the three scaffolded constructs used in this study.

**Extended Data Figure 5. Cryo-EM structure determination of Tet_P6b_ + Zika Virus xrRNA.** The Cryo-EM processing pipeline in cryoSPARC for the Tet_P6b_ + Zika Virus xrRNA construct.

**Extended Data Figure 6. Tet_P6b_ + Zika Virus xrRNA principal component analysis.** Component 0 of a three-component principal component analysis of the Tet_P6b_ + Zika Virus xrRNA map. Maps from frame 1, 3, and 5 show local motion in the P4 helix of the Zika Virus xrRNA. This component largely encompasses a flexing motions of stem P6b in the scaffold and the appended domain as a result.

**Extended Data Figure 7. Comparison of Tet_P6b_ + Zika Virus xrRNA with crystal structures of other xrRNAs.** Models of the Tet_P6b_ + Zika Virus xrRNA (**A**), Zika Virus xrRNA (**B**), and Murray Valley encephalitis virus (MVEV) xrRNA (**C**). **D.** Superimposition of the three xrRNAs.

**Extended Data Figure 8. Cryo-EM structure determination of Tet_P6b_ + TABV xrRNA.** The Cryo-EM processing pipeline in cryoSPARC for the Tet_P6b_ + TABV xrRNA construct.

**Extended Data Figure 9. Cryo-EM structure determination of Tet_P6b_ + *T. petrophila* Fluoride riboswitch.** The Cryo-EM processing pipeline in cryoSPARC for the Tet_P6b_ + *T. petrophila* Fluoride riboswitch construct.

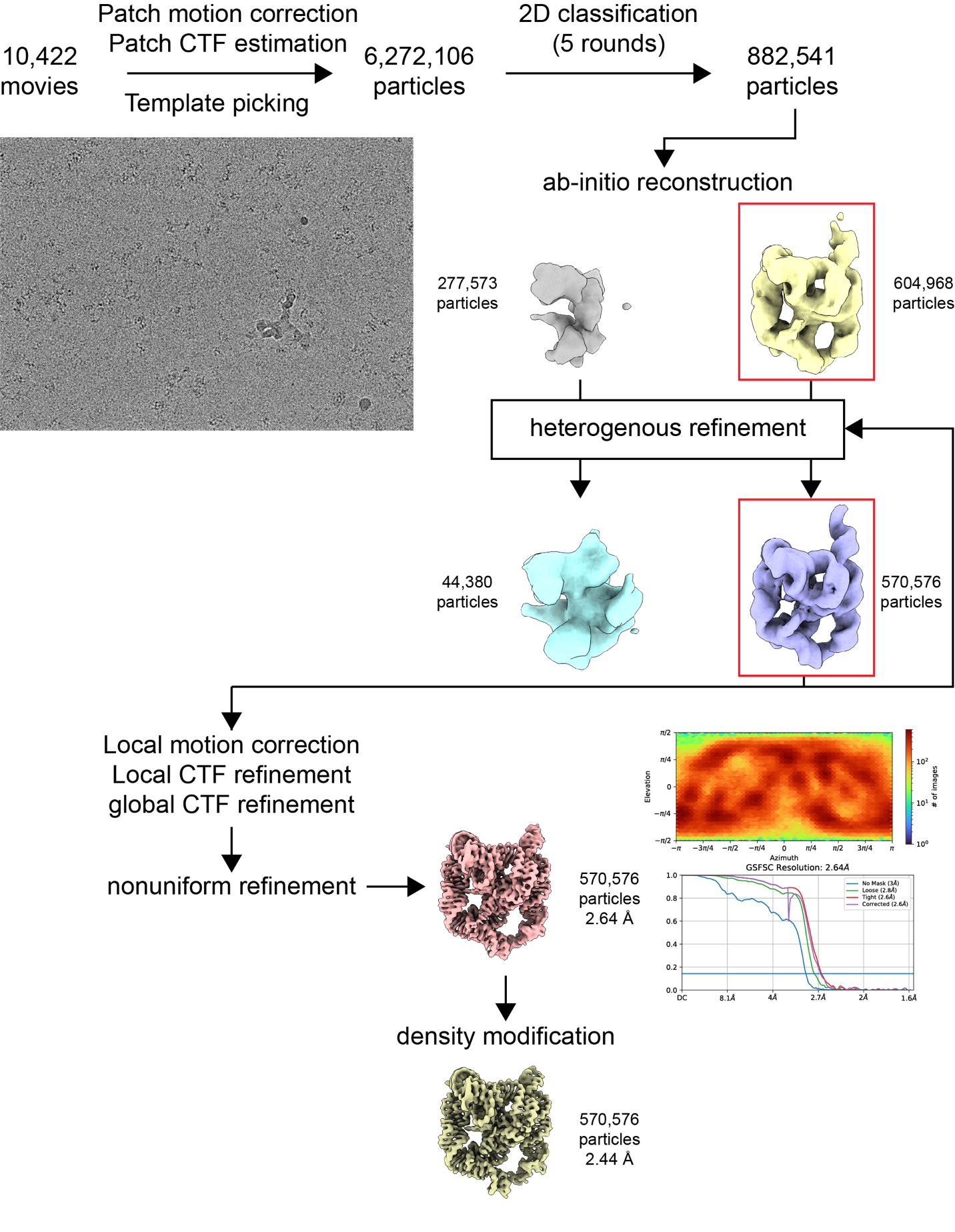

**Extended Data Figure 1. Cryo-EM structure determination of Tet_P6b_.**

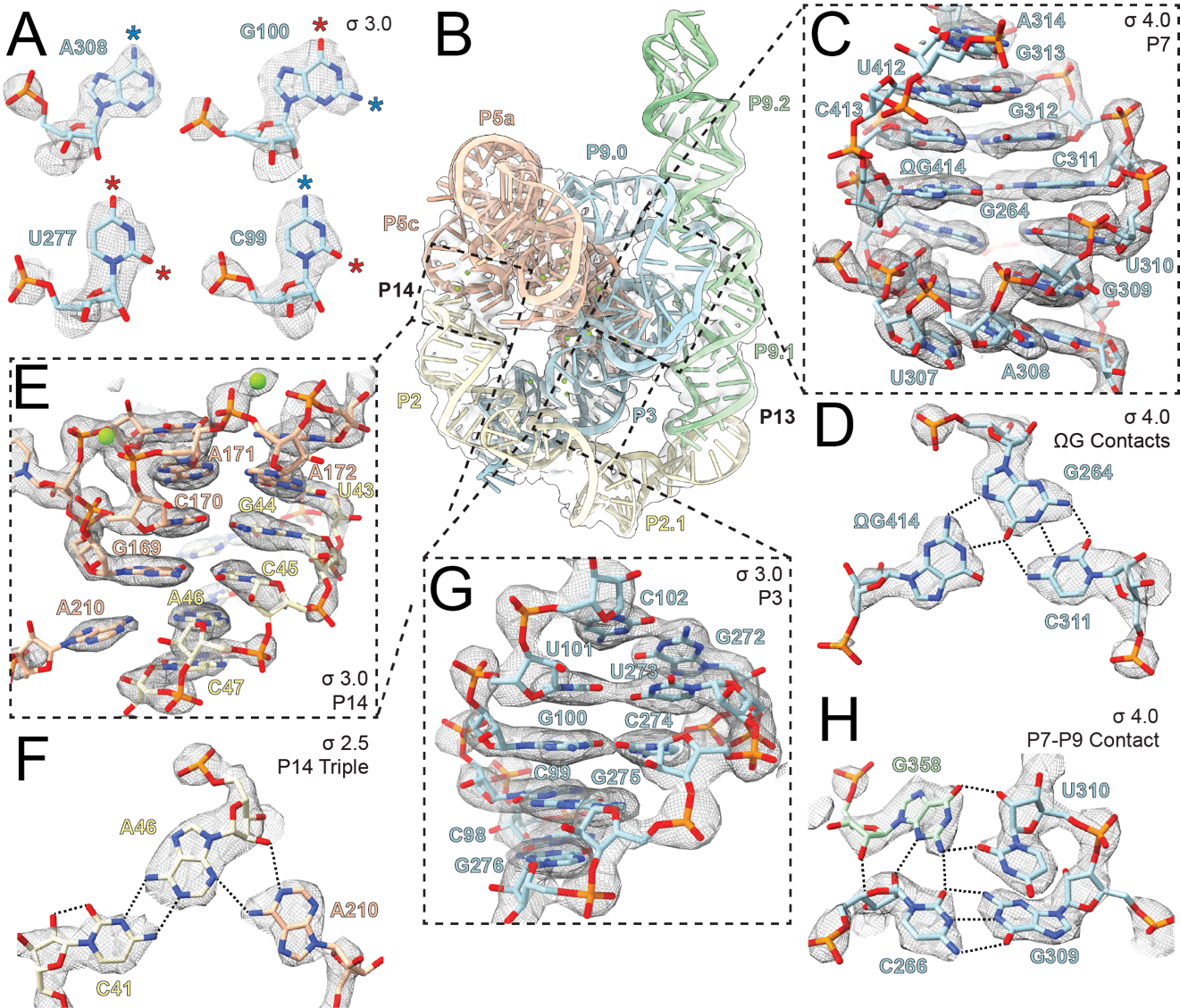

**Extended Data Figure 2. Tet_P6b_ reconstruction details.**

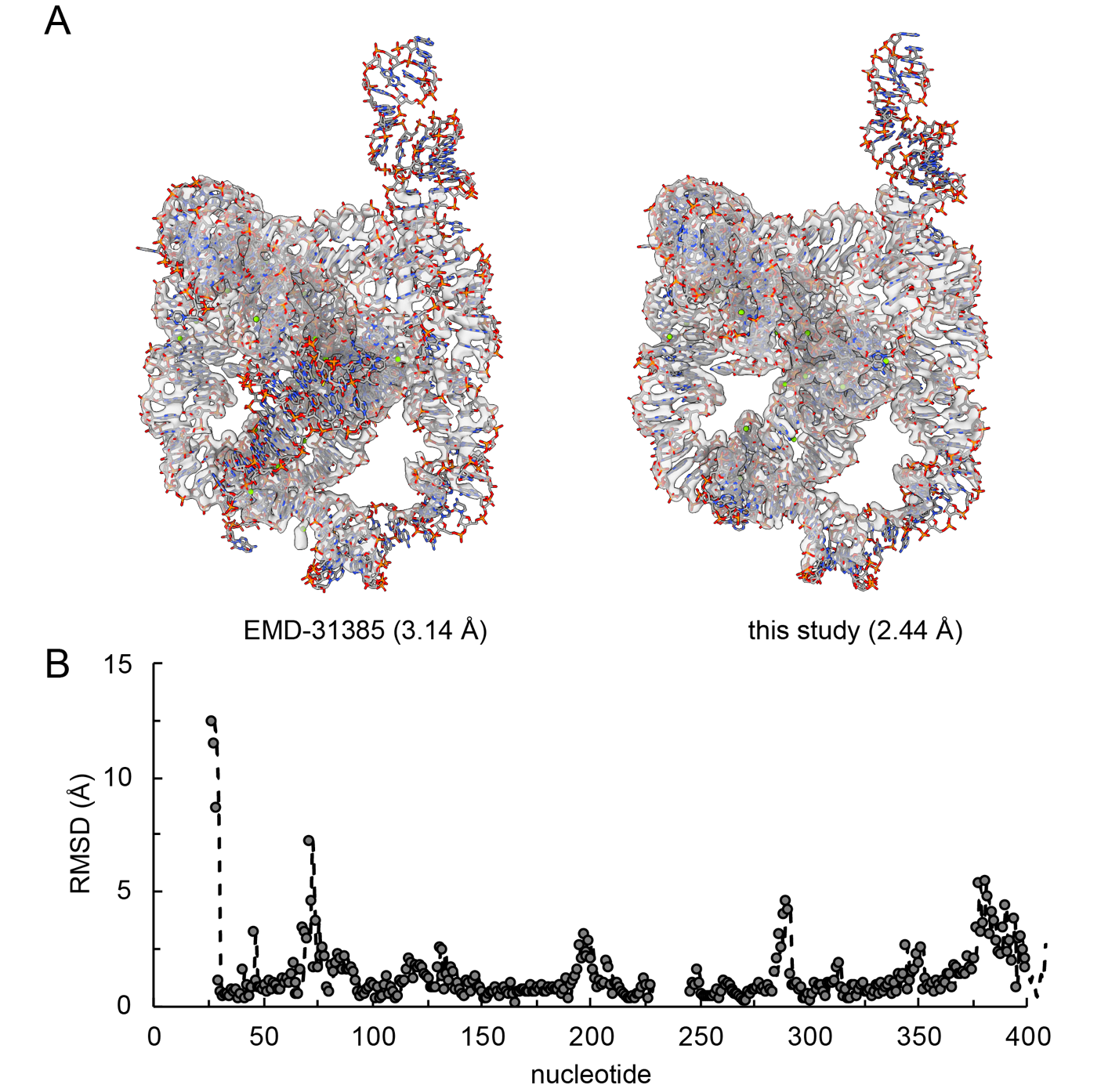

**Extended Data Figure 3. Comparison of Tet_P6b_ to a published tetrahymena ribozyme structure.**

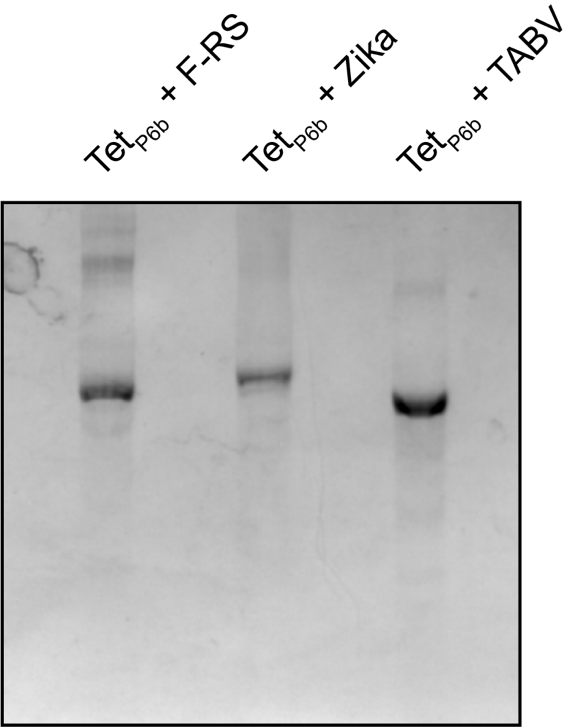

**Extended Data Figure 4. Native gel analysis of scaffolded RNAs.**

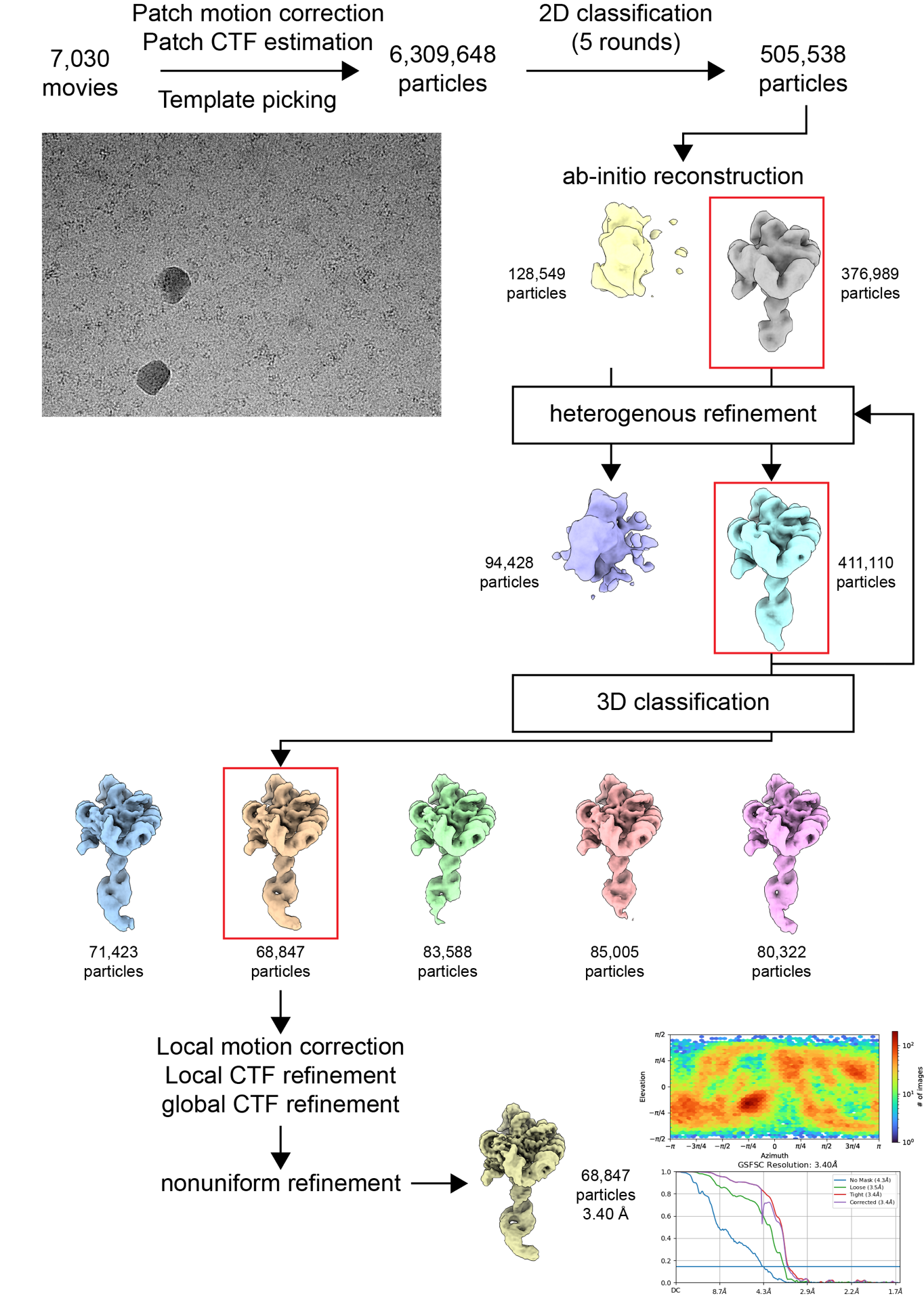

**Extended Data Figure 5. Cryo-EM structure determination of Tet_P6b_ + Zika Virus xrRNA.**

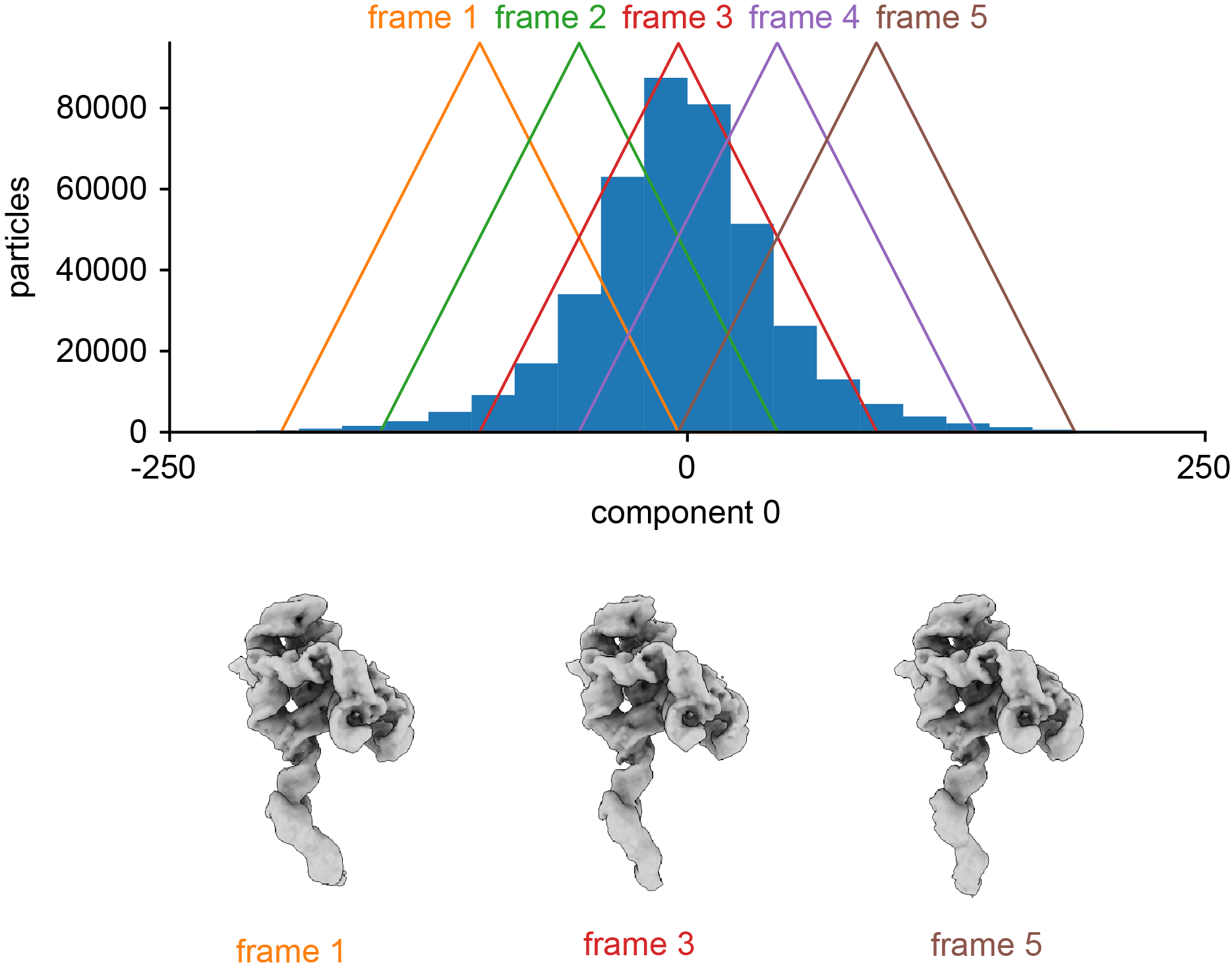

**Extended Data Figure 6 Tet_P6b_ + Zika Virus xrRNA principal component analysis.**

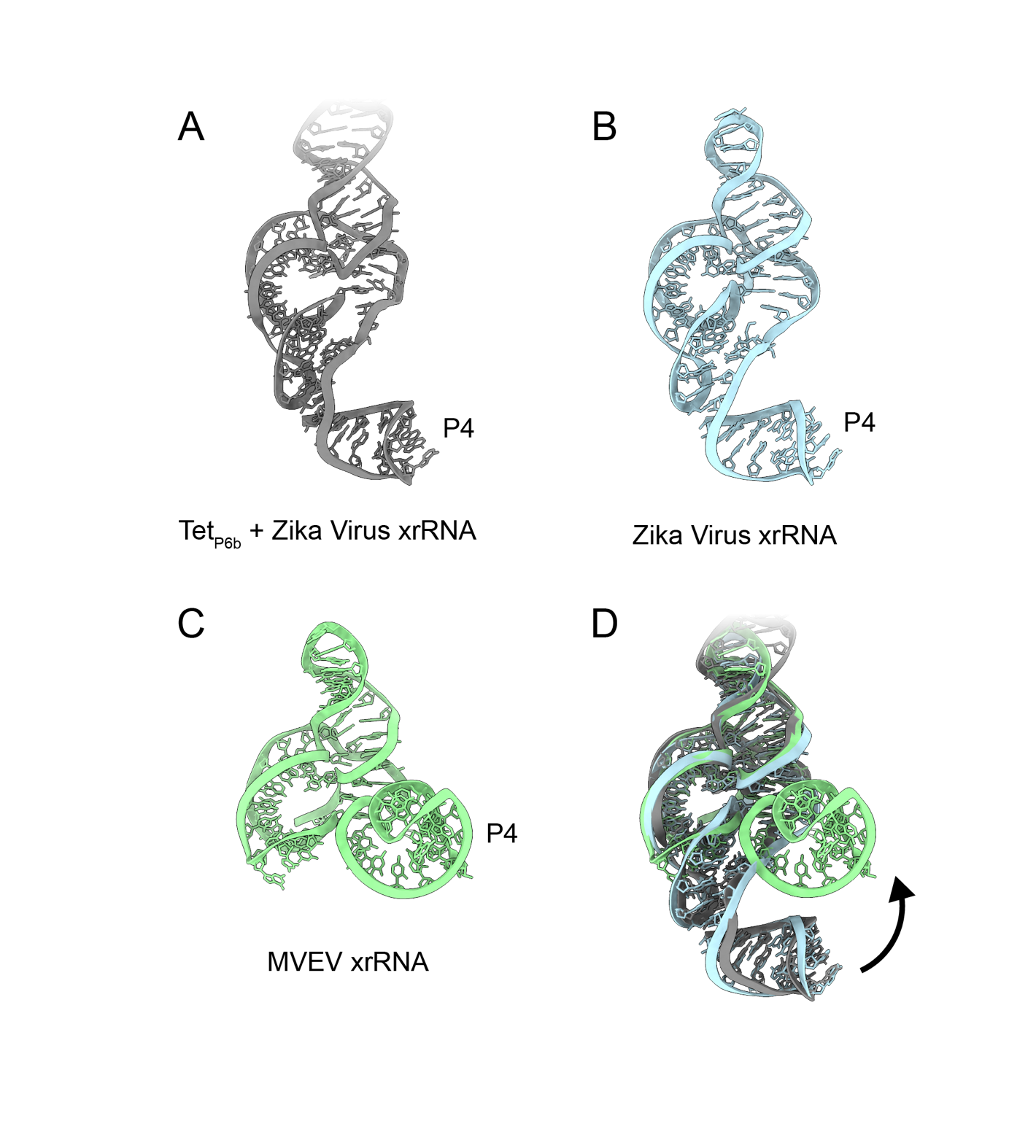
 **Extended Data Figure 7. Comparison of Tet_P6b_ + Zika Virus xrRNA with crystal structures of other xrRNAs.**

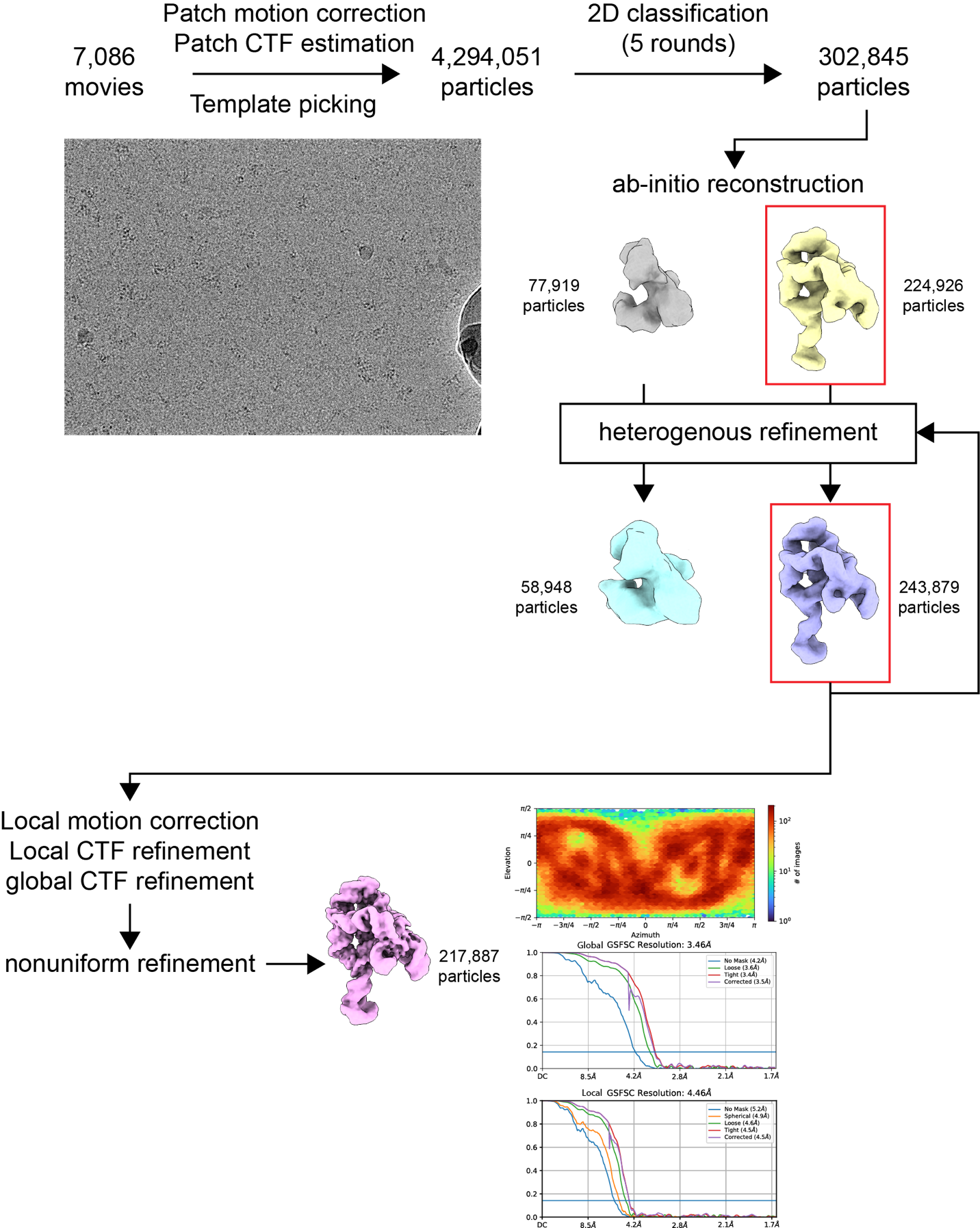

**Extended Data Figure 8. Cryo-EM structure determination of Tet_P6b_ + TABV xrRNA.**

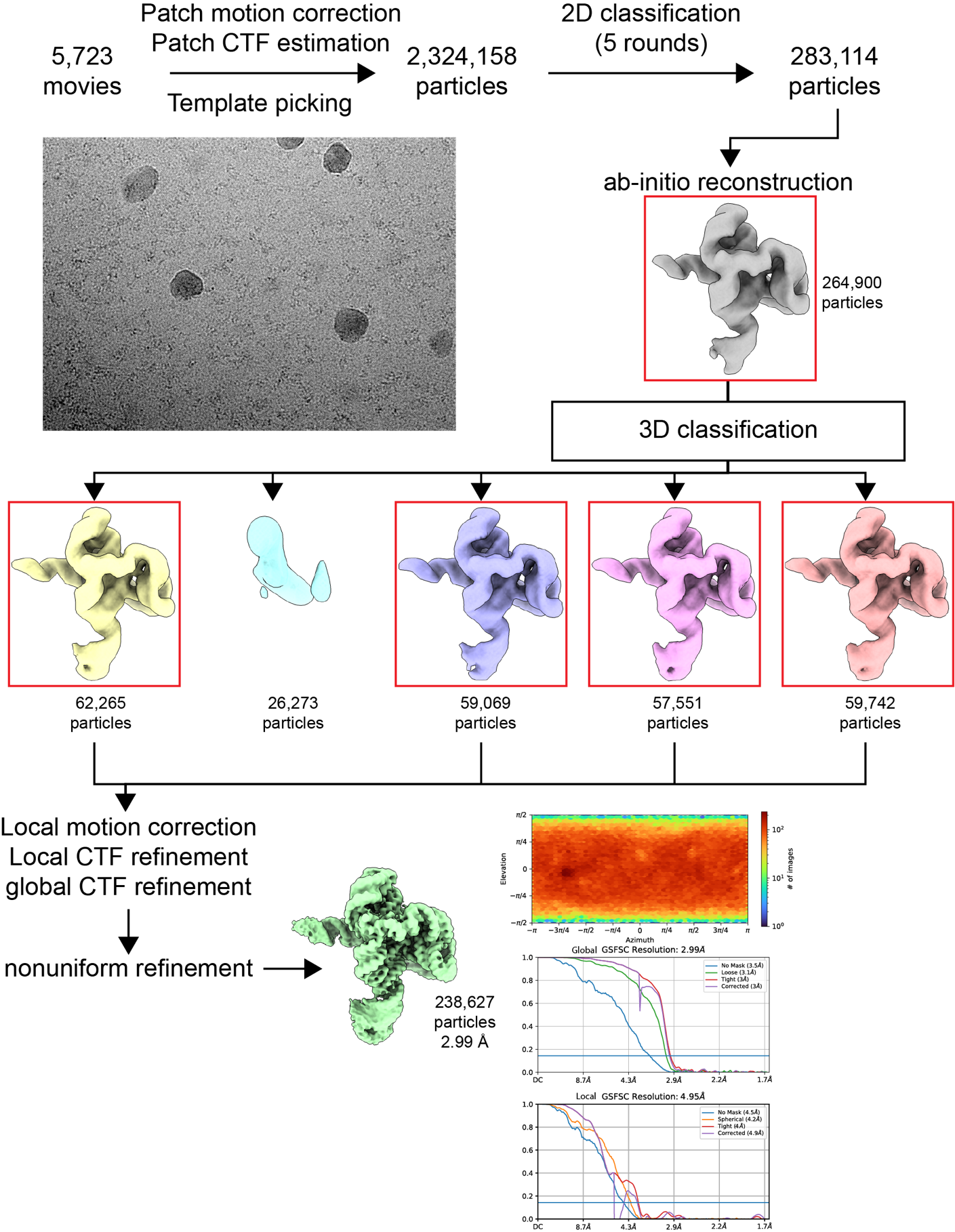

**Extended Data Figure 9. Cryo-EM structure determination of Tet_P6b_ + *T. petrophila* Fluoride riboswitch.**
