## Supplemental Material Tables for "A Generalizable Scaffold-Based Approach for Structure Determination of RNAs by Cryo-EM"

**Contents:**

**Supplementary Table S1:** Cryo-EM data collection, refinement, and validation statistics

**Supplementary Table S2:** Oligos used in this study

|  | TetP6b | TetP6b + Zika xrRNA | TetP6b + TABV xrRNA | TetP6b + Fluoride riboswitch |
| --- | --- | --- | --- | --- |
|  | (EMDB-xxxx) | (EMDB-xxxx) | (EMDB-xxxx) | (EMDB-xxxx) |
|  | (PDB xxxx) | (PDB xxxx) | (PDB xxxx) | (PDB xxxx) |
| **Data collection and processing** |  |  |  |  |
| Grid Type | C-flat 1.2/1.3 | C-flat 1.2/1.3 | C-flat 1.2/1.3 | C-flat 1.2/1.3 |
| Microscope | FEI Titan Krios | FEI Titan Krios | FEI Titan Krios | FEI Titan Krios |
| Camera | K3 | K3 | K3 | K3 |
| Magnification | 105,000 | 105,000 | 105,000 | 105,000 |
| Voltage (kV) | 300 | 300 | 300 | 300 |
| Electron exposure (e–/Å^2^) | 50 | 50 | 50 | 50 |
| Defocus range (μm) | -0.8 to -2.0 | -0.8 to -2.1 | -0.8 to -2.2 | -0.8 to -2.3 |
| Pixel size (Å) | 0.8464 | 0.8464 | 0.8464 | 0.8464 |
| Symmetry imposed | C1 | C2 | C3 | C4 |
| Initial particle images (no.) | 6,272,106 | 6,309,648 | 4,294,051 | 2,324,158 |
| Final particle images (no.) | 570,576 | 68,847, | 217,887 | 238,627 |
| Map resolution (Å) | 2.44 | 3.40 | 3.46 | 2.99 |
| FSC threshold | 0.143 | 0.143 | 0.143 | 0.143 |
| Map resolution range (Å) | 2.340-6.669 | 2.919-7.061 | 2.981-7.904 | 2.520-7.156 |
| **Refinement** |  |  |  |  |
| Initial model used (PDB code) | 7EZ2 | 7EZ2, 5TPY | 7EZ2, 7K16 | 7EZ2, 4ENC |
| Model resolution (Å) | 2.44 | 3.40 | 3.46 | 2.99 |
| FSC threshold | 0.143 | 0.143 | 0.143 | 0.143 |
| Mask CC | 0.880 | 0.860 | 0.854 | 0.868 |
| Model resolution range (Å) | 2.0-6.0 | 3.0-8.0 | 3.0-8.0 | 2.5-8.0 |
| Map sharpening *B* factor (Å^2^) | -76.3 | -85.8 | -156.0 | -85.1 |
| Model composition |  |  |  |  |
| Non-hydrogen atoms | 7750 | 9454 | 8929 | 8959 |
| RNA bases | 361 | 440 | 415 | 417 |
| Ligands | 27 | 28 | 27 | 32 |
| *B* factors (Å^2^) |  |  |  |  |
| RNA | 70.22 | 150.16 | 62.7 | 62.68 |
| Ligand | 32.83 | 76.48 | 18.94 | 28.32 |
| R.m.s. deviations |  |  |  |  |
| Bond lengths (Å) | 0.003 | 0.021 | 0.011 | 0.006 |
| Bond angles (°) | 0.559 | 1.321 | 0.947 | 0.915 |
| Validation |  |  |  |  |
| MolProbity score | 2.03 | 3.32 | 2.38 | 2.21 |
| Clashscore | 1.46 | 49.52 | 4.55 | 2.75 |
| Poor rotamers (%) | 0 | 0 | 0 | 0 |

**Supplementary Table 1. Cryo-EM data collection, refinement, and validation statistics**

| **Sequence 5' to 3'** | GCTAACGGATCCTAATACGACTCACTATAGGGTGTGTCTTGGATCGCGCGGTGATATGGATGCAGTTCACAGACTAAATGTCGGTCGGGGAAGATGTATTCTTCTCATAAGATATAGTCGGACCTCTCCTTAATGGGAGCTAGCGGATGAAGTGATGCAACACTGGAGCCGCTGGGAACTAATTTGTATGCGAAAGTATATTGATTAGTTTTGGAGTACTCGTTGGAGGGAAAAGTTATCAGGCATGCACCTGGTAGCTAGTCTTTAAACCAATAGATTGCATCGGTTTAAAAGGCAAGACCGTCAAATTGCGGGAAAGGGGTCAACAGCCGTTCAGTACCAAGTCTCAGGGGAAACTTTGAGATGGCCTTGCAAAGGGTATGGTAATAAGCTGACGGACATGGTCCTAACCACGCAGCCAAGTCCTAAGTCACCGCAGGCACGTAATAAAGCGAGGGGTTCGAATCCCCCCGTTACCCCCGGTAGGGGCCCATCTAGAGCATAC | GCTAACGGATCCTAATACGACTCACTATA**GGGTCAGGCCGGC**TGATATGGATGCAGTTCACAGACTAAATGTCGGTCGGGGAAGATGTATTCTTCTCATAAGATATAGTCGGACCTCTCCTTAATGGGAGCTAGCGGATGAAGTGATGCAACACTGGAGCCGCTGGGAACTAATTTGTATGCGAAAGTATATTGATTAGTTTTGGAGTACTCGTTGGAGGGAAAAGTTATCAGGCATGCACCTGGTAGCTAGTCTTTAAACCAATAGATTGCATCGGTTTAAAAGGCAAGACCGTCAAATTGCGGGAAAGGGGTCAACAGCCGTTCAGTACCAAGTCTCAGGGGAAACTTTGAGATGGCCTTGCAAAGGGTATGGTAATAAGCTGACGGACATGGTCCTAACCACGCAGCCAAGTCCTAAGTCA**GTCGCCACAGTTTGGGGAAAGCTGTGCAGCCTGTAACCCCCCCACGAAAGTGGG**TCTAGAGCATAC | GCTAACGGATCCTAATACGACTCACTATA**GGCAAGGTACGGC**ATATGGATGCAGTTCACAGACTAAATGTCGGTCGGGGAAGATGTATTCTTCTCATAAGATATAGTCGGACCTCTCCTTAATGGGAGCTAGCGGATGAAGTGATGCAACACTGGAGCCGCTGGGAACTAATTTGTATGCGAAAGTATATTGATTAGTTTTGGAGTACTCGTTGGAGGGAAAAGTTATCAGGCATGCACCTGGTAGCTAGTCTTTAAACCAATAGATTGCATCGGTTTAAAAGGCAAGACCGTCAAATTGCGGGAAAGGGGTCAACAGCCGTTCAGTACCAAGTCTCAGGGGAAACTTTGAGATGGCCTTGCAAAGGGTATGGTAATAAGCTGACGGACATGGTCCTAACCACGCAGCCAAGTCCTAAGT**GCCGTAGGGGCTTGAGAACCCCCCCTCCCCACTC**TCTAGAGCATAC | GCGAGTGAATTCTAATACGACTCACTATA**GGGCGATGAGGCCCGCCCAAACTGCCCG**ATATGGATGCAGTTCACAGACTAAATGTCGGTCGGGGAAGATGTATTCTTCTCATAAGATATAGTCGGACCTCTCCTTAATGGGAGCTAGCGGATGAAGTGATGCAACACTGGAGCCGCTGGGAACTAATTTGTATGCGAAAGTATATTGATTAGTTTTGGAGTACTCATTGGAGGGAAAAGTTATCAGGCATGCACCTGGTAGCTAGTCTTTAAACCAATAGATTGCATCGGTTTAAAAGGCAAGACCGTCAAATTGCGGGAAAGGGGTCAACAGCCGTTCAGTACCAAGTCTCAGGGGAAACTTTGAGATGGCCTTGCAAAGGGTATGGTAATAAGCTGACGGACATGGTCCTAACCACGCAGCCAAGTCCTAAGTC**GGGCTGATGGCCTCTACTG** | GCTAACGGATCCTAATACGACTCACTATAGGG | TGGGCCCCTACCGGGGGTAACG | TAATACGACTCACTATAGGGTCAGGCCG | CCCACTTTCGTGGGGGGG | TAATACGACTCACTATAGGCAAGGTACGGC | GAGTGGGGAGGGGGGG | TAATACGACTCACTATAGGG | CAGTAGAGGCCATCAGCCCG |
| --- | --- | --- | --- | --- | --- | --- | --- | --- | --- | --- | --- | --- |
| **Oligo** | TetP6b gBlock | TetP6b + Zika Virus xrRNA gBlock | TetP6b + TABV xrRNA gBlock | TetP6b + Fluoride riboswitch gBlock | TetP6b forward primer | TetP6b reverse primer | TetP6b + Zika Virus xrRNA forward primer | TetP6b + Zika Virus xrRNA reverse primer | TetP6b + TABV xrRNA forward primer | TetP6b + TABV xrRNA reverse primer | TetP6b + Fluoride riboswitch forward primer | TetP6b + Fluoride riboswitch reverse primer |

**Supplementary Table 2. Oligos used in this study**
